# Re-inventing pathogen passage for social microbes

**DOI:** 10.1101/2022.08.04.502769

**Authors:** Tatiana Dimitriu, Wided Souissi, Peter Morwool, Alistair Darby, Neil Crickmore, Ben Raymond

## Abstract

Passage experiments that sequentially infect hosts with parasites have long been used to manipulate virulence. However, in many invertebrate pathogens passage has been applied naively without a full theoretical understanding of how best to select for increased virulence. This has led to very mixed results. Understanding the evolution of virulence is complex because selection on parasites occurs across multiple spatial scales with potentially different conflicts operating on parasites with different life-histories. For example, in social microbes, strong selection on replication rate within hosts can lead to cheating and loss of virulence, because investment in public goods virulence reduces replication rate. In contrast, selection acting at a between host scale maintains virulence by selecting on parasite population size. In this study we tested how different scales of selection and varying mutation supply affect evolution of virulence against resistant hosts in the specialist insect pathogen *Bacillus thuringiensis*., aiming to optimize methods for strain improvement against a difficult to kill insect target. We show that selection for infectivity using competition between sub-populations in a metapopulation prevents social cheating, acts to retain key virulence plasmids and facilitates increased virulence. Increased virulence was associated with reduced efficiency of sporulation, and loss of function in putative regulatory genes but not with altered expression of known virulence factors. Selection in a metapopulation provides a broadly applicable tool for improving the efficacy of biocontrol agents. Moreover, a structured host population can facilitate artificial selection on infectivity, while selection on life history traits such as faster replication or larger population sizes can reduce virulence can reduce virulence in social microbes.

## Introduction

Passage, the repeated infection and re-isolation of a microbe in a host, has been used as a tool for the manipulation of parasite virulence for decades, as well as a means of testing evolution of virulence theory (Ebert 1998; Raymond & Erdos 2022). Passage has been successfully used as a means of produced attenuated live vaccines: sequential infection of animal tissue cultures can lead to loss of virulence in human hosts and has been used to produce polio and yellow fever vaccines among others (Sabin & Boulger 1973; Barrett 2017). Conversely, in some circumstances adaptation of viruses to a particular host via passage can lead to increased virulence (Ebert 1998). Although there are complex relationships between virulence and fitness in natural populations (Frank 1996; Gandon, Jansen, & van Baalen 2001; Alizon, Hurford, Mideo, & Van Baalen 2009), artificial inoculation means that many of the negative consequences of high virulence in terms of reducing opportunities for transmission will not operate in laboratory conditions and so it may be possible to increase the virulence of naturally occurring pathogens via artificial selection.

Increasing pathogen virulence via passage has long been a goal in biocontrol research as researchers have sought to manipulate the efficacy and / or host range of microbes that are potential biocontrol agents (Raymond & Erdos 2022). Naïve passage designs, in which the aim is simple infection and re-infection, without any other selection pressure or modifications have proved successful for a number of baculoviruses (Maleki-Milani 1978; Kolodny-Hirsch & Van Beek 1997; Berling et al. 2009). The success of the simple passage of baculoviruses can be partly explained by the high genetic diversity found in natural populations (Shapiro, Lynn, & Dougherty 1992; Thézé et al. 2014). However, another reason naïve passage can be effective is if reproductive rate within hosts is positively correlated with virulence, as is assumed by many classical models of virulence (Frank 1996; Gandon, Jansen, & van Baalen 2001; Alizon, Hurford, Mideo, & Van Baalen 2009). Even in the simplest passage design there will be competition between genetic variants within host, and we would expect that this within host competition would favour higher replication rate, ignoring issues such as antagonism and evasion or manipulation of immunity (Massey, Buckling, & Ffrench-Constant 2004; Raberg et al. 2006). This means that simple isolation and re-infection can select for the genotypes which grow fastest within hosts with the net result of an increase in virulence without the need for any additional selection pressure.

Nevertheless, research on the social biology of micro-organisms emphasizes that increased replication rate does not necessarily lead to increased virulence (Buckling & Brockhurst 2008; Raymond & Erdos 2022). Many bacterial pathogens, for instance, invest considerable resource in producing virulence factors such as toxins or siderophores that are important for infecting hosts or accessing host resources such as iron (West & Buckling 2003; Raymond, West, Griffin, & Bonsall 2012; Diard, Garcia, et al. 2014). Investing in these resources can slow replication rates, this means that intense within-host competition can select for cheaters which can out-compete virulent genotypes in mixed infections, although cheaters are expected to have low fitness in single strain infections. In addition, increased genetic diversity within infections (low relatedness) will also tend to favour fast-replicating genotypes such as cheaters. These social biology concepts are relevant both for clinically important microbes and for biocontrol agents. Experimental evolution with entomopathogenic nematodes, for instance, shows increasing opportunities for cheating can lead to attenuation and extinction, and may explain historical problems with unstable virulence in laboratory culture of these parasites (Shapiro-Ilan & Raymond 2016).

In this study we aimed to apply social evolution theory to increase the virulence of the Gram-positive invertebrate pathogen, *Bacillus thuringiensis* (Bt). Bt is a valuable model for testing novel passage regimes partly because its virulence factors are extremely well characterized (Adang, Crickmore, & Jurat-Fuentes 2014) and because its social biology is well-studied (Raymond et al. 2012; Zhou et al. 2014; Cornforth, Matthews, Brown, & Raymond 2015; van Leeuwen, Neill, Matthews, & Raymond 2015). Bt obligately requires pore-forming toxins for infection. These are produced at sporulation in the cadaver but solubilized in the midgut after ingestion; the toxins disintegrate the midgut epithelium and allow these microbes access to the haemocoel (Adang et al. 2014). In some insects hosts the production of quorum-regulated phospholipases and cellulases complement the action of the δ-endotoxins and facilitate host invasion (Salamitou et al. 2000; Zhou et al. 2014). A range of other toxins eg Vegetative Insecticidal Proteins (Vips) are produced by most Bt strains, although their contribution to pathogenicity is less clear (Chakroun et al. 2016). While the entomocidal toxins of Bt vary considerably in structure and potency, most have high selective potency against insect pests (Palma et al. 2014) this is the main reason why Bt dominates the microbial biocontrol market and supplies the vast majority of insecticidal toxins for genetically-modified crops (Bravo, Likitvivatanavong, Gill, & Soberon 2011).

There is considerable interest in identifying mutants or proteins that can overcome host resistance in Bt (Badran et al. 2016). From a theoretical point of view, it may also make sense to target passage experiments at partially resistant hosts. Based on fitness landscape theory, we expected that rapid, experimentally tractable evolution would be more likely in a pathogen of low fitness rather than one already at an adaptive peak and easily capable of infecting hosts (Poelwijk, Kiviet, Weinreich, & Tans 2007). We conducted experimental evolution using a host species commonly targeted by Bt products: the diamondback moth *Plutella xylostella*, selecting a well characterized genotype with a high level of resistance to *B. thuringiensis kurstaki*. For social microbes, the selection pressure acting to maintain virulence does not come from within host competition for rapid growth-it has to come from competition between populations in terms of total population size (yield) (Griffin, West, & Buckling 2004) or number of hosts infected (Raymond et al. 2012). This is because the fitness benefits (or public goods) provided by cooperation in known social pathogens either increase the efficient use of host resource or, in the case of δ-endotoxins, provide access to host tissues.

In this study, we tested passage regimes that would maximize selection based on yield (population size within hosts) or infectivity. Selecting for yield involved combining all cadavers in a replicate into a single inoculum pool at each passage so that pathogens with higher yield are better represented in the next round of infections (Griffin et al. 2004; Shapiro-Ilan & Raymond 2016) (Fig 1). This type of between-host competition has previously produced modest increases in virulence in Bt (Garbutt, Bonsall, Wright, & Raymond 2011). Pooling subpopulations also mimics a high pathogen dispersal rate within natural populations (Kümmerli, Gardner, West, & Griffin 2009). The second selection regime involved directly selecting for infectivity. To do this we imposed competition between sub-populations within a metapopulation (each metapopulation constituting an independent replicate), so that only the 50% most infectious subpopulations are used to initiate the next round of infection. The most infectious subpopulations are divided and initiate two new pathogen subpopulations in the next round of infection. The infectivity selection treatment therefore uses a form of budding dispersal and so may have additional benefits in terms of preserving population structure (Gardner & West 2006; Kümmerli et al. 2009). It should be emphasized that these selection treatments are not exclusive or perfect, in other words we cannot prevent some selection for infectivity in the yield treatment (infections have to happen) nor some selection for yield in the infectivity treatment (bacteria have to grow in cadavers) but we are attempting to maximize different types of selection with our experimental treatments.

**Figure 1.**
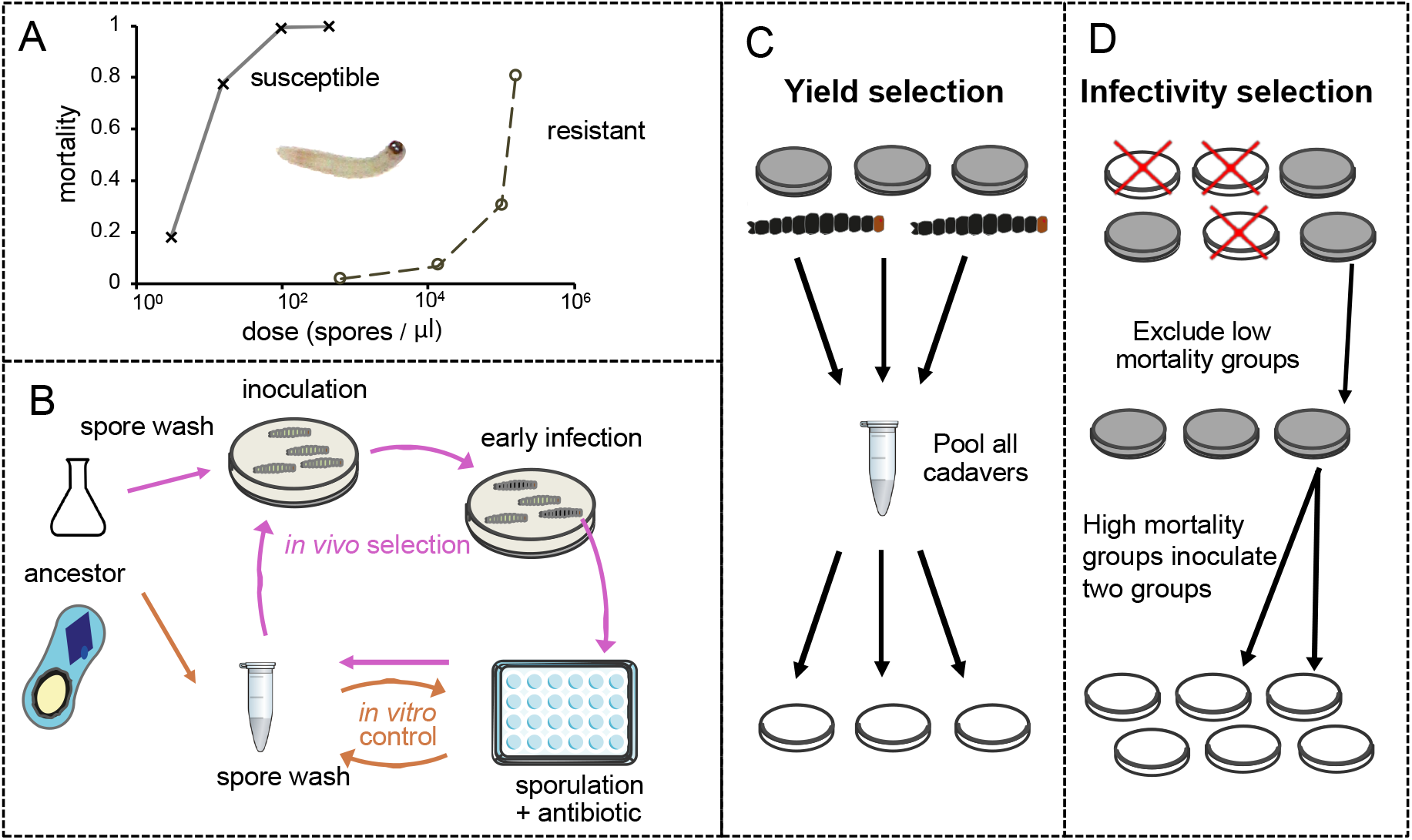
Experimental design to increase virulence of *Bacillus thuringiensis* against resistant insects. **(A)** Bioassays using the Bt ancestor strain 7.1.o show substantial differences in mortality between our focal Bt resistant insect VLFR and its near-isogenic Bt susceptible counterpart VLSS (*F*_*1,5*_ *=* 34.1, *P* = 0.0021). The LC50 of the susceptible insects was estimated at 7.4 CFU μl^-1^ (95% confidence limits 6.9-7.9 CFU μl^-1^), while the LC50 of the VL FR resistant is 5×10^5^ CFU μl^-1^ (Table S1). Selection experiments began with a dose that would kill 20-30% of insects. (**B)** Selection cycles, performed separately for strains with mutator or wild type mutation rates. For *in vivo* selection treatments, cells selected from killed larvae were inoculated into sporulation medium. *In vitro* growth, sporulation and purification steps were common to the *in vivo* selection and *in vitro* control. To prevent the invasion of cheaters not contributing to cooperative virulence, two *in vivo* selection treatments were compared. (**C)** In the yield selection treatment, all cadavers from a replicate lineage were pooled and inoculated together for spore production. **(D)** Under group selection half of the sub-populations, those causing the lowest mortality, are terminated at the end of each round of selection, while spores from the remaining sub-populations are divided and used to infect two-subpopulations in the next round of infection.

Finally, we tested how increasing the mutation supply would affect virulence evolution, a common approach in directed evolution (Selifonova, Valle, & Schellenberger 2001). Bacterial clones with elevated mutation rates and defective proof-reading genes, or mutators, are prevalent in pathogenic species (Taddei et al. 1997), and we increased mutation supply by approximately 25-fold by carrying out selection in a mutator, made by disruption of the *mutS* gene, as well as in the wild type ancestor. At the end of passage experiments we bioassayed for changes in virulence in evolved lineages (as well as clones isolated from those lineages and assayed for changes in life history (spore production, competitive fitness) and tested whether social conflicts might have affected experimental outcomes. Finally, in order to explore the possible mechanisms for observed increased in virulence in evolved clones we explored if changes in the expression of known virulence factors could explain results and conducted whole genome sequencing on a selection of evolved clones.

## Methods

### Insects

The Cry1Ac resistant line “VL-FR” used for experiments was previously derived from a cross of NOQA-GE with the diet adapted susceptible line Vero Beach, this line was selected for resistance to Cry1Ac as described previously (Zhou et al. 2018). We established a population that was fixed for resistance using a PCR screen of resistance alleles in the parental population. An insect strain with similar genetic background (also derived from a NOQA-GE X Vero Beach cross) was established from F2 offspring but in this case fixed for susceptibility to Cry1Ac, this line was denoted ‘VL-SS’ (Zhou et al. 2018).

### Bacteria, plasmids and construction of strains for this work

The ancestor *B. t. kurstaki* strain was obtained by transforming the original field isolate of the 7.1.o strain (Raymond et al. 2009) with pHT315-gfp plasmid containing the *ermB* gene conferring erythromycin resistance (Zhou et al. 2014). It was maintained *in vitro* with 10μg/mL erythromycin. When grown from insect cadavers, 30μg/mL erythromycin was used to inhibit growth of Gram-negative bacteria.

The mutator (mut) strain was obtained by inactivating the *mutS* gene in 7.1.o via disruption. The *cat* gene conferring chloramphenicol resistance, was digested from the pAB2 plasmid using *KpnI* (Bravo, Agaisse, Salamitou, & Lereclus 1996), this fragment was inserted at position 1439 of *mutS* coding sequence. The sequence between positions 601 to 1837 of *mutS* gene coding sequence with the *cat* insert was synthesized in pUC57 from GenScript after adding *BamHI* sites on both ends. This synthesized *BamHI* fragment was cloned into the thermosensitive vector pRN1501 (Lereclus et al. 1992). The resulting plasmid was introduced into 7.1.o strain, and clones with recombination of the interrupted *mutS* sequence into the chromosome were obtained after growth at 42°C by screening for chloramphenicol (Chl) resistance and loss of erythromycin resistance (Lereclus et al. 1992). Insertion was confirmed by PCR and sequencing of *mutS* locus. The mut strain was grown in the presence of 5μg/ml Chl.

The mutation rate of the ancestor with wild type *mutS* (wt) and mut strain was calculated with fluctuation assays. 32 independent cultures per strain were inoculated into HCO medium containing respectively erythromycin 300 μg/ml for the wt strain and Chl 5 μg/ml for the mut strain, by 10^7^-fold dilution from LB overnight cultures. After 24h at 30°C, cultures were plated on LB agar plates containing 100 μg/ml rifampicin. Mutation rates and confidence intervals were calculated with the Ma-Sandri-Sarkar (MSS)-maximum likelihood method using FALCOR (Hall, Ma, Liang, & Singh 2009). Wt mutation rate towards rifampicin resistance was 7×10^−10^ [95%CI 4.2×10^−10^ to 1.02×10^−9^) and mut strain mutation rate was 1.76×10^−8^ [95% CI 1.48×10^−8^ to 2.08×10^−8^], showing a 25-fold increase compared to the wt strain. The mutator, prior to passage, did not show any change in virulence relative to the ancestor (log_10_ LC50 = 4.9, test for difference from ancestor-effect size 0.21, SE = 1.22, *p* = 0.87). The competitive fitness of the mutator strain under the conditions of the passage experiment also did not differ from the wt ancestor (*t* = 0.49, *p* = 0.63, means ±SE of mut and wt are 1.09 ±0.06 and 1.13 ±0.06 respectively).

### Passage experiment

Diamondback moth larvae used for selection were reared on autoclaved artificial diet (Baxter et al. 2011) without supplementary antibiotics until 3^rd^ instar. *Bacillus thuringiensis* strains used for the selection experiment were first grown from single colonies into 100 ml of sporulating media (HCO) with appropriate antibiotic, then pasteurized (65°C, 20min). Selection was then performed with 4 replicate lineages per treatment.

For infection, sterile diet was cut into quarters in 55ml dishes, and each quarter was inoculated with 90μl of spore suspension containing 5×10^4^ to 10^5^ spores/μl (chosen to produce 25-30% mortality over 2-4 days for resistant diamondback moth) and dried. Ten 3^rd^ instar larvae were then added per 55ml dish. Insect death was recorded daily from two days after infection. When total mortality reached 25-30%, the earliest cadavers were transferred to micro-centrifuge tubes with sterile toothpicks. After two days at room temperature, 0.85% NaCl solution (saline) was added to the tubes. Cadavers were homogenized with pellet pestles, then the tubes were briefly vortexed and the suspension was transferred to wells containing 1 ml of sporulation medium in 24-well growth plates, for spore amplification *in vitro*. Sporulation medium was HCO (Lecadet, Blondel, & Ribier 1980) containing polymyxin (100 IU/ml) (Oxoid) and an additional relevant antibiotic: erythromycin 300 μg/ml for wt lineages containing pHT315-gfp or chloramphenicol 5 μg/ml for mut lineages containing the chromosomal *cat* marker. Rows of inoculated wells from the same line of selection were alternated with empty rows to prevent and control for contamination between lines. The plates were then sealed with tape to limit evaporation and placed at 30°C with 150rpm agitation for three days.

Selection of cadavers and inoculation into sporulation medium depended on experimental treatment (Figure 1). For the group selection treatment, each lineage was split into 12 populations maintained separately. The 6 populations with highest larval mortality were selected; 6 other populations were discarded. If the same mortality was observed for several populations, the ones with earliest observed deaths were selected. A maximum of two-three cadavers from selected populations were transferred into separate tubes (6 total). 50 μl saline was added in each tube, and the suspension from each tube was inoculated into separate wells. After spore purification (described below), spores from each well were used to inoculate two populations in the next selection round. For the pooled selection treatment, each lineage was split into 6 populations, which were pooled together at each round of selection. Cadavers from all 6 populations in the pooled treatment were transferred to the same microfuge tube. 300 μl saline was then added and the suspension was distributed into 6 wells for inoculation. For replicate lineages with more than 30% death across populations at time of selection, the number of cadavers taken from populations with most deaths was reduced in order to select 30% cadavers overall and maintain comparable population sizes across treatments at inoculation.

After three days of growth, 200 μl spore aliquots were transferred to 96-well PCR plates with aluminium sealing film for quantification, infection in next passage and storage. Plates were centrifuged at 3000 g for 15 min, the supernatant was removed and spores were resuspended in 100 μl 0.85% NaCl. Plates containing spores to be used in next infection step were pasteurized (20 min, 65°C) and kept at 10°C until use, the others were stored at - 20°C. Spore density was estimated by diluting 10 μl of spores into 90 μl saline and measuring OD600nm in a 96-well microplate reader. A subset of the samples was plated at appropriate dilutions on LB-agar (LBA) plates, in order to visually check for contamination and calibrate OD600nm measurements.

The *in vitro* control selection treatment followed the same protocol without infection of insects and was similarly performed on 4 lineages per strain with 5 rounds of selection. Approximately 5×10^4^ spores were used to inoculate 1ml sporulation medium (containing antibiotics at the same concentration used for *in vivo* selection). After 3 days of growth, spores were centrifuged in 1.5ml micro-centrifuge tubes and resuspended in 250 μl 0.85% NaCl, then pasteurized and kept at 10°C until use.

### Phenotypic and genotypic characterization of evolved bacteria

*Bioassays*. After five rounds of selection, the evolved lineages were frozen at -80°C (LB 20% glycerol) without pasteurization. These frozen stocks were used to inoculate sporulation medium (HCO) for assays of whole lineages, as above but without antibiotics. Clones were isolated from population samples by streaking single colonies three times successively on LB agar plates supplemented with 10 μg/ml erythromycin (wt clones) or 5 μg /ml chloramphenicol (mut clones). Clone nomenclature in figures and tables indicates genetic background (W for wild type mutation rate, M for mutator), selection treatment (y for yield, i for infectivity selected, v for *in vitro* control) with a letter for independent lineage and a number for clone, *e*.*g*. Wia1, wild type, infectivity selected, lineage a, clone 1.

For bioassays, mixed populations or clones were inoculated into 1 ml HCO (without antibiotics) and grown for 3 days at 30°C, with 150 rpm agitation. Spore concentration was measured by plating on LBA after pasteurization. Lineage-level assays were done twice with independent spore stocks. Each assay was performed with at least three doses with 60 insects per dose (between 10^4^ and 1.2×10^5^ spores μl^-1^ for *B. t. kurstaki* in resistant *P. xylostella*). Initial screens of clone level variation examined 6 clones per lineage; these used 15-30 insects per dose at two doses. Clonal screens were analyzed in terms of viable spores (counted as colony forming units on LBA) but also in terms of dilution factor of inocula relative to purified spore stocks as some clones displayed variable germination rates on LBA. Clones of interest were further characterized in additional bioassays, with methods that followed lineage assays, these assays were repeated at least twice.

The virulence of each independent lineage was assayed by scoring proportional mortality (Figure 2A and B) after 3 days (early mortality) and 5 days (late mortality). This analysis used data from two independent bioassays using independently grown spore stocks (minimizing effects due to variation in growth / amplification *in vitro*). Bioassays used three doses, and analyses reported in Figure 2A used mortality data that were normalized against the ancestor: the average mortality (across the three doses) of each evolved lineage replicate was divided by the average mortality measured in the ancestor within the same assay.

**Figure 2.**
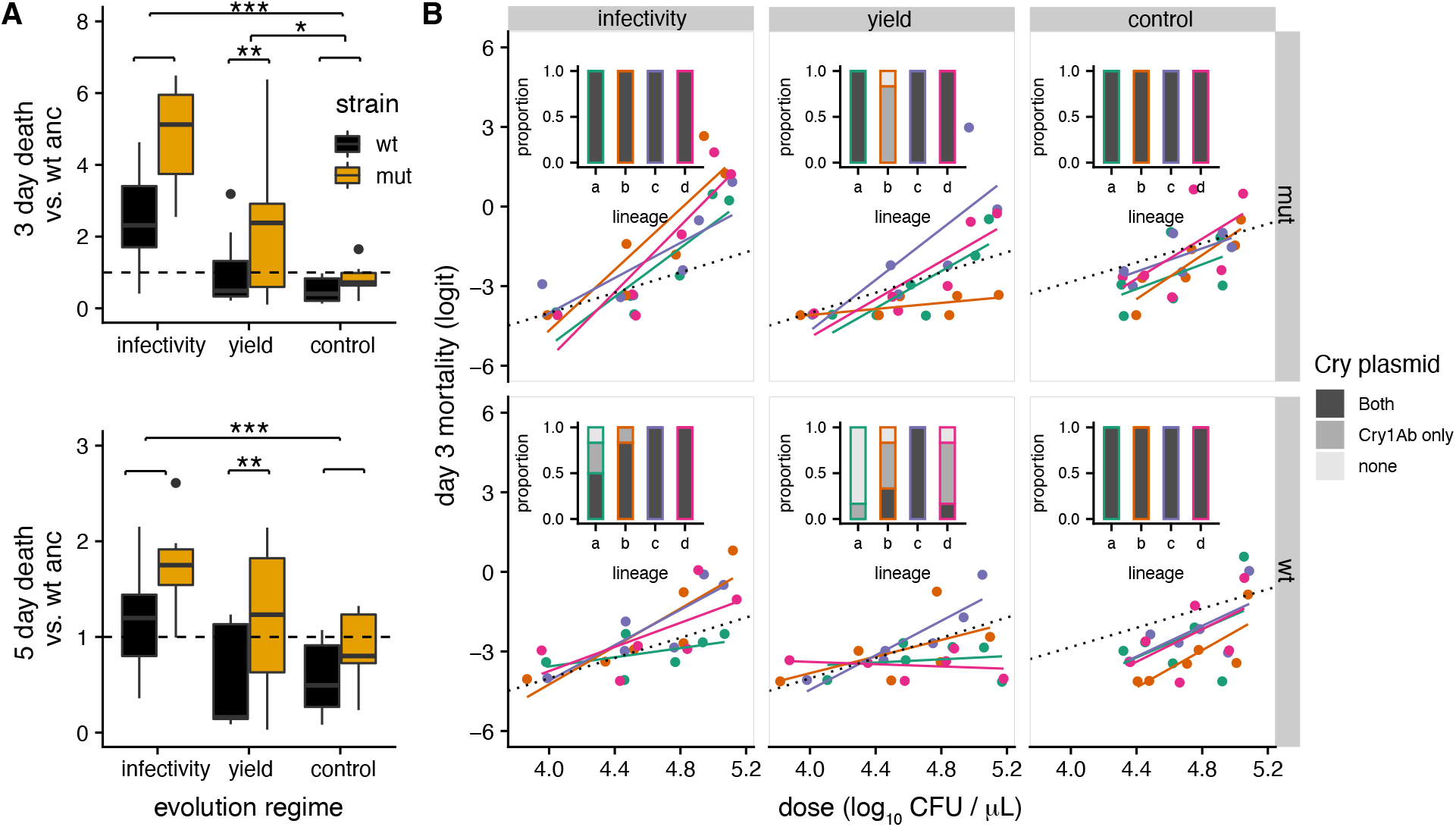
Mutation rate and group selection shape evolution of virulence and retention of public good virulence factors (Cry toxins). Data shown are from two separate bioassays performed with independent spore preparations of evolved lineages after five rounds of selection (**A)** Proportional mortality (relative to the ancestor) at 3 and 5 days after inoculation for different treatments, control refers to *in vitro* passaged controls. The centre value of the boxplots shows the median, boxes the first and third quartile, and whiskers represent 1.5 times the interquartile range; outliers are shown as dots (N=8, 4 lineages per treatment in two assays). Both mutation rate (*F*_1,44_ = 12.3, *P* = 0.001 at 3 days, *F*_1,44_ = 10.6, *P* = 0.002 at 5 days) and selection level (3 days - *F*_2,44_ = 21.4, *P* < 0.001, 5 days - *F*_2,44_ = 8, *P* = 0.0011) affected virulence. *Post hoc* tests confirm increased virulence for infectivity versus yield selected treatments and for mutators versus wild type lineages (Tukey tests, all *P* < 0.01). (**B)** Logit-transformed mortality 3 days after inoculation is shown as a function of Bt dose. For reference the dotted black line shows the dose response for the ancestor, the replicate lineages within each treatment are colour coded. Insets show for each lineage the proportion of clones containing no Cry toxins (light grey), Cry1Ab only (dark grey) or both Cry1A and Cry2A (black) toxin genes (N=6 clones per lineage). See also Table S1 and Figures S1-S2 for results of clone-level virulence assays.

### Genetic analysis of Cry toxin gene complements

The ancestor *B. t. kurstaki* 7.1.o strain bears genes for several expressed toxins (Cry1Ac, Cry2Aa, Cry1Aa) on a large plasmid (homologous to CP009999, 317kb) and one additional toxin gene (Cry1Ab) on a smaller plasmid (homologous to CP010003, 69kb)(Tables S2-3), this smaller plasmid is also homologous to the pHT73 plasmid described in the *Bt* strain HD-73 (Méric et al. 2018). Loss of each of those plasmids was detected in several clones with whole-genome sequencing. We extended those results by estimating Cry toxin plasmid loss for all clones from evolved line by PCR (6 clones from each insect evolved lineage, and 2 clones from each *in vitro* lineage). Primers were designed and validated with the sequenced clones. Primers Cry2-F and Cry2-R amplify a 1.1kb product from the large toxin-encoding plasmid (CP009999). Primers Cry1-F and Cry1-R amplify a 1.4kb product when any Cry1A gene is present; no amplification corresponds to loss of both *cry* gene bearing plasmids.

PCR used HotStar Taq (Qiagen) with cycling parameters: 5 min at 95°C, 30 × (45 s at 94°C, 1 min at 50°C [Cry2 primers] or 53°C [Cry1 primers], 90 s at 72°C), 10 min at 72°C. To provide DNA templates, fresh colonies were resuspended into 100μl dH_2_O, frozen 20 min at -80°C then boiled 10 min. 5μl supernatant was used in 20μl PCR mix. To exclude false negative results, each negative result was repeated 3 times from fresh colonies.

### Whole-genome sequencing

The unmarked ancestral *B. t. kurstaki* 7.1.o strain was sequenced with PacBio, data were produced using SMRTbell® Express Template Prep Kit 2.0 following the manufacturer’s recommendations. This resulted in a 15-20 kb insert library which was sequencing on a PacBio Sequel system using one cell per library and 10-hour sequencing movie time. The data was processed to provide CCS and single pass data and assembled using Unicycler version 0.4.7 (Wick, Judd, Gorrie, & Holt 2017).

For insect-evolved lineages, the two clones with highest virulence based on a preliminary screening were chosen. At least two clones from each endpoint evolved lineage, in addition to the wt ancestor (7.1.o wt pHT315-gfp) and the mutator (7.1.o ΔmutS) were sequenced. DNA was extracted using Qiagen DNeasy Blood and Tissue genomic extraction kits, after appropriate lysis for Gram-positive bacteria. Sequencing was performed on an Illumina HiSeq 2500 at a minimum of 30x coverage with 125bp paired end sequencing.

All Illumina reads were trimmed with Trimmomatic (Bolger, Lohse, & Usadel 2014). Unicycler version 0.4.7 was used to perform a hybrid assembly using PacBio self-corrected reads and Illumina reads (Wick et al. 2017). This combination of read types resulted a final assembly that included small plasmids that were otherwise lost from the PacBio data only assemblies, due to DNA fragment size section been larger than the size of the plasmids. Short read data were used to validate and aid the comparison of plasmid genomes between the reference strain HD-1 and the 7.1.o strain (this study) by mapping the short reads to the assembly (Li 2012) (Tables S2-S3).

The final assembly was then checked against other *Bacillus thuringiensis kurstaki* stains using MAUVE (Darling, Tritt, Eisen, & Facciotti 2011) to check for possible genome errors. The assembled genome was then run though the software PROKKA (Seemann 2014) providing a draft annotation. The PROKKA flag --usegenus –genus *Bacillus* was used to improve taxon specific gene annotations. The final assembly was then checked against other Illumina whole genome shotgun data from the selection experiments were then analysed with snippy version 4.4.0 (https://github.com/tseemann/snippy) to align reads against the newly assembled reference and used to call single nucleotide polymorphisms (SNPs). The Snippy SNP data was used to generate a Jukes Cantor Neighbor-Joining (1000 boot strap replicates) phylogeny of isolates, implemented with Geneious Prime 2022.2.1.

### Fitness measurements

Measurements of relative fitness used a mutant of the 7.1.o isolate constructed by transformation with a pHT315-rfp plasmid containing the *tetR* gene conferring tetracycline resistance (Zhou et al. 2014). This version of the ancestor therefore has a plasmid with a similar backbone to the evolved wild type strains but is distinguishable on the basis of its distinct antibiotic resistance. Relative fitness was calculated from an estimate of relative reproductive rates (Malthusian parameter) of two genotypes (Zhou, Slamti, Lereclus, & Raymond 2020).

Fitness was measured in conditions covering a cycle of the experimental design used for selection *in vivo*. A 50/50 mix of spores of the two competitors was used for insect infection. The protocol then followed the experimental design used for selection *in vivo*, except insect cadavers were processed individually and transferred to different wells for sporulation. Moreover, the sporulation medium contained no antibiotics. After spore cleaning, mixes were plated on antibiotic-free medium; tetracycline 10ng/μl (for the standard competitor ancestor) and plates without antibiotics. The density of evolved clones in cadavers was calculated by subtracting competitor counts from antibiotic-free plates total counts, as the *cat* gene insertion proved unreliable at allowing growth on chloramphenicol and we wished to use a common method for both wt and mut clones.

### Assessment of toxin production

Bt strains were cultured in HCO medium at 30°C with shaking (200rpm) hours in the presence/absence of erythromycin (5μg/ml) as required. Sporulation and crystal production were monitored using phase contrast microscopy. For the assessment of crystal protein production 1ml of the culture was centrifuged and the pellet resuspended in 1ml of water. Two rounds of sonication were performed to break open un-lysed cells before the final pellet was resuspended in 1ml of water. Serial dilutions were plated out in order to calculate the concentration of colony forming units (CFUs). SDS-PAGE analysis was then performed on equal numbers of CFUs to compare production of crystal proteins. For the assessment of Vip3A production, culture supernatant was passed through a 0.22 μm cellulose acetate filter and to 1 ml of the filtrate 50μl of 2% sodium deoxycholate was added and incubated for 30 min on ice. 150 μl of 100% Trichloroacetic acid was then added, the sample was vortexed and incubated overnight at 4°C. The precipitate was spun down at 15000g for 10 min, the supernatant removed, and the pellet was washed twice with ice cold 70% acetone then air dried. Following the addition of 20 μl water, the mixture was sonicated for 2 min to resuspend the pellet. For immunodetection of Vip3Aa samples were boiled with SDS sample buffer and spotted onto a nitrocellulose membrane. A Vip3A antibody (a kind gift from Professor Juan Ferré) was used in association with an anti-rabbit IgG HRP-linked antibody for ECL detection.

### Statistical analysis

All statistical analyses were carried out in R (v4.0.4) (https://www.R-project.org). Bioassay and mortality data were analyzed using one of two methods. For comparisons between independently evolved lineages we used generalized linear modelling (glms) with logit link functions and quasibinomial errors to correct for overdispersion. However, to test for treatment effects, or account for random effects associated with different lineages within treatments, we used generalized linear mixed models (glmer) in the package *lme4* (Bates, Machler, Bolker, & Walker 2015). Although we attempted to use glmer models with binomial errors, these commonly failed to converge, especially in more complex models with dose X treatment interactions. Instead, we used the conservative approach of arc-sine transforming proportional mortality and using normal errors. The glm and glmer approaches produce qualitatively similar results. Hypothesis testing in glmers used likelihood ratio tests after model simplification. LC50s and their standard errors were calculated using the *dose*.*p* function in the *MASS* package (Venables & Ripley 2002) after fitting simple logit models using log_10_ dose and quasibinomial errors.

Analyses of spore yield also used glmers with lineage fitted as a random effect, while the analysis of competitive fitness used evolved clone as a random effect and genotype (mutator or wild type) and number of toxins as explanatory variables. Model assumptions (normality, heteroscedascity) were checked with graphical analyses and qq plots.

## Results

### Changes in virulence

Endpoint assays, after five rounds of selection (*ca*. 60 generations), showed that increased mutation rate and infectivity selection between populations resulted in increased virulence relative to both the ancestor strain and the *in vitro* controls (Fig 2A). Differences between treatments were larger when assessing early mortality, suggesting changes in virulence affected timing and overall levels of mortality (Fig 2A). Clonal assays of late mortality show that mutation rate and infectivity selection increased virulence via interactions with dose (mixed effect *glm* with replicate as random effect: dose*scale of selection interaction *df* = 3, LR = 16.95, *p* < 0.001; dose*strain interaction *df* = 1, LR = 12.59, *p* < 0.001; Fig S1, S2). We identified a number of clones with substantial increases in virulence, quantified as a reduction of LC50 of more than an order of magnitude (Table S1) from endpoint assays.

### Fitness, cheating and patterns of Cry toxin plasmid carriage

Cry toxins are encoded on two plasmids in Bt *kurstaki -* a mega-plasmid containing Cry1Aa, Cry1Ac and Cry2Aa, and a circa80kb plasmid carrying Cry1Ab (Méric et al. 2018). We used PCR primers to screen for putative cheater mutants that had lost these plasmids during selection. We observed numerous events of toxin gene loss in clones isolated from *in vivo* evolved lineages (Fig 2B bar plots). Importantly, the infectivity selection regime was more effective at retaining these plasmids and preventing invasion of putative cheaters (Fig 2B, Fisher’s exact test two-tailed *p* = 0.00032). The division of successful subpopulations via budding dispersal in our infectivity selection treatment has the potential to reduce social conflict by maintaining high relatedness and by purging the population of subpopulations dominated by cheats (Kümmerli et al. 2009). In order to test whether mutants that had lost Cry-toxin plasmids were cheaters, we measured the fitness of one clone from each lineage in competition experiments that used a marked derivate of the ancestor. Since mutants carrying fewer Cry toxin genes had higher competitive fitness (mixed model likelihood ratio test (LRT) *df* = 1, Likelihood ratio (LR) = 9.05, *p* < 0.0026) and lower virulence (*glm* F_1, 64_ = 67.1, *p* < 0.0001, Fig 3B) these data were consistent with the hypothesis that low virulence mutants were free-loading on investment in Cry toxins by other strains in their lineages (Fig 3A,B) (Raymond et al. 2012).

**Figure 3.**
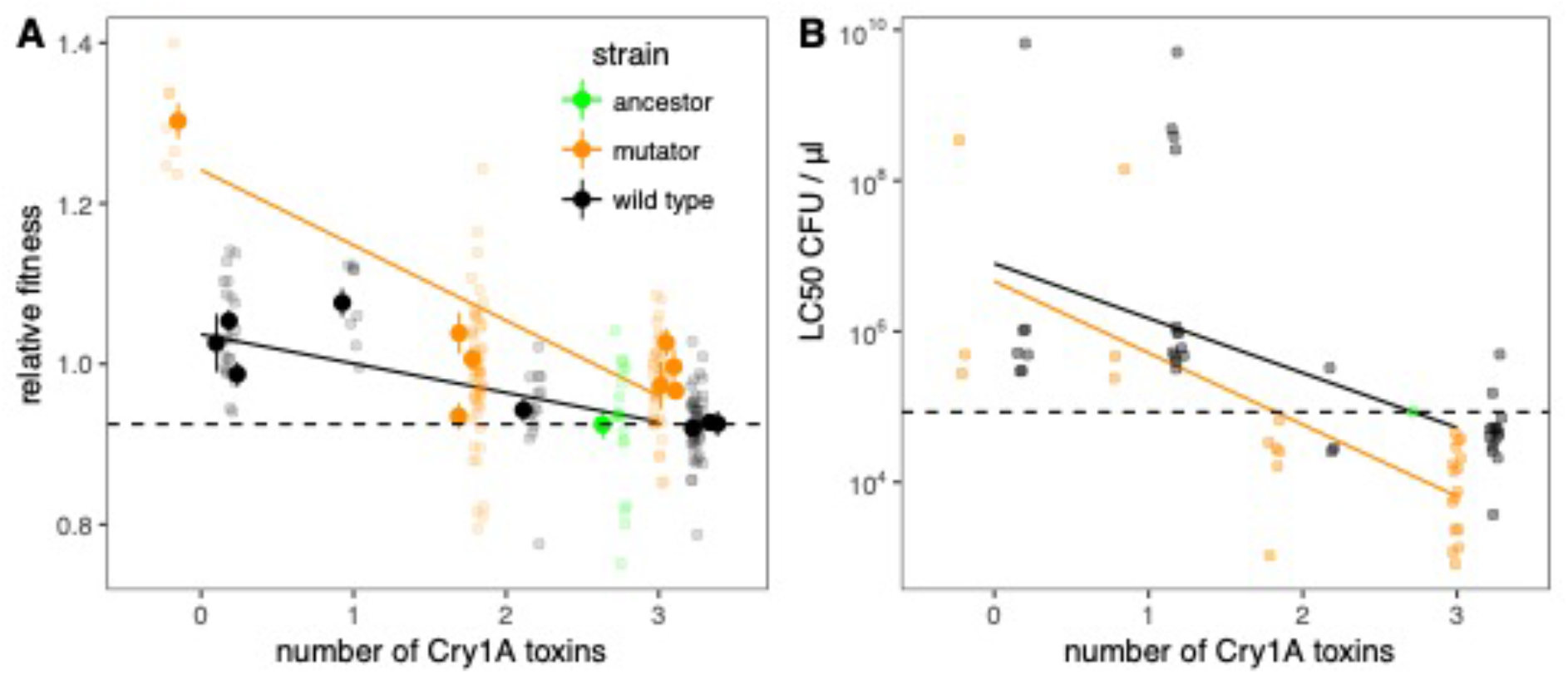
Social cheating and fitness in experimentally evolved Bt lines. Evolved clones (n = 16) that had lost virulence plasmids encoding Cry genes had higher competitive fitness (A) and lower virulence (B) than the ancestor. LC50s are shown for all evolved clones with toxin complement data (n = 70). Means±SE of fitness for each clone are plotted as solid points, raw data are plotted at 50% transparency, the dashed line is a reference line for the ancestor while solid lines show fitted statistical models (*glms* with genotype and number of Cry toxins).

Mutator lineages retained more plasmids than wild type lineages (Fig 2B; Fisher’s exact test two-tailed *p* = 0.006). After accounting for toxin production, mutators evolved greater competitive fitness than wild type lineages (Fig 3A; mixed model, *df* = 1, LR = 7.5, *p =* 0.006) and had increased virulence (*i*.*e*. lower LC50s ; *glm* F_1, 63_ = 7.35, *p* < 0.0157, Fig 3B) suggesting that increasing mutation rate increased the supply of beneficial alleles without reducing overall fitness. Since the original mutator mutant has indistinguishable fitness from the wild type ancestor (see Star Methods), initial genetic background does not explain this pattern.

### Spore production

The other major life-history that we explored was the efficiency of spore production in the sporulation media used in the selection experiments. We saw lower spore production in the mutator and the infectivity-selected lineages, the treatments that produced higher virulence (Fig 4A, mixed models, selection treatment *df* = 3, LR = 11.3, *p* = 0.01, mutation rate *df* = 1, LR = 8.36, *p* < 0.01). Importantly, between lineages lower spore production was associated with higher virulence (Fig 4B, *F*_*1,14*_ *=* 10.5, *p* < 0.01, adj. *R*^*2*^ = 0.39). This is biologically significant because toxin production is linked to sporulation in Bt, especially the Cry1A toxins that are the dominant virulence factors in our focal strain (Deng, Peng, Song, & Lereclus 2014). In addition, we selected for sporulation by heat-treating preparations at the end of each round of selection to kill vegetative cells and possible contaminants. This suggests that changes in virulence are unlikely to be caused by Cry1A toxins.

**Figure 4.**
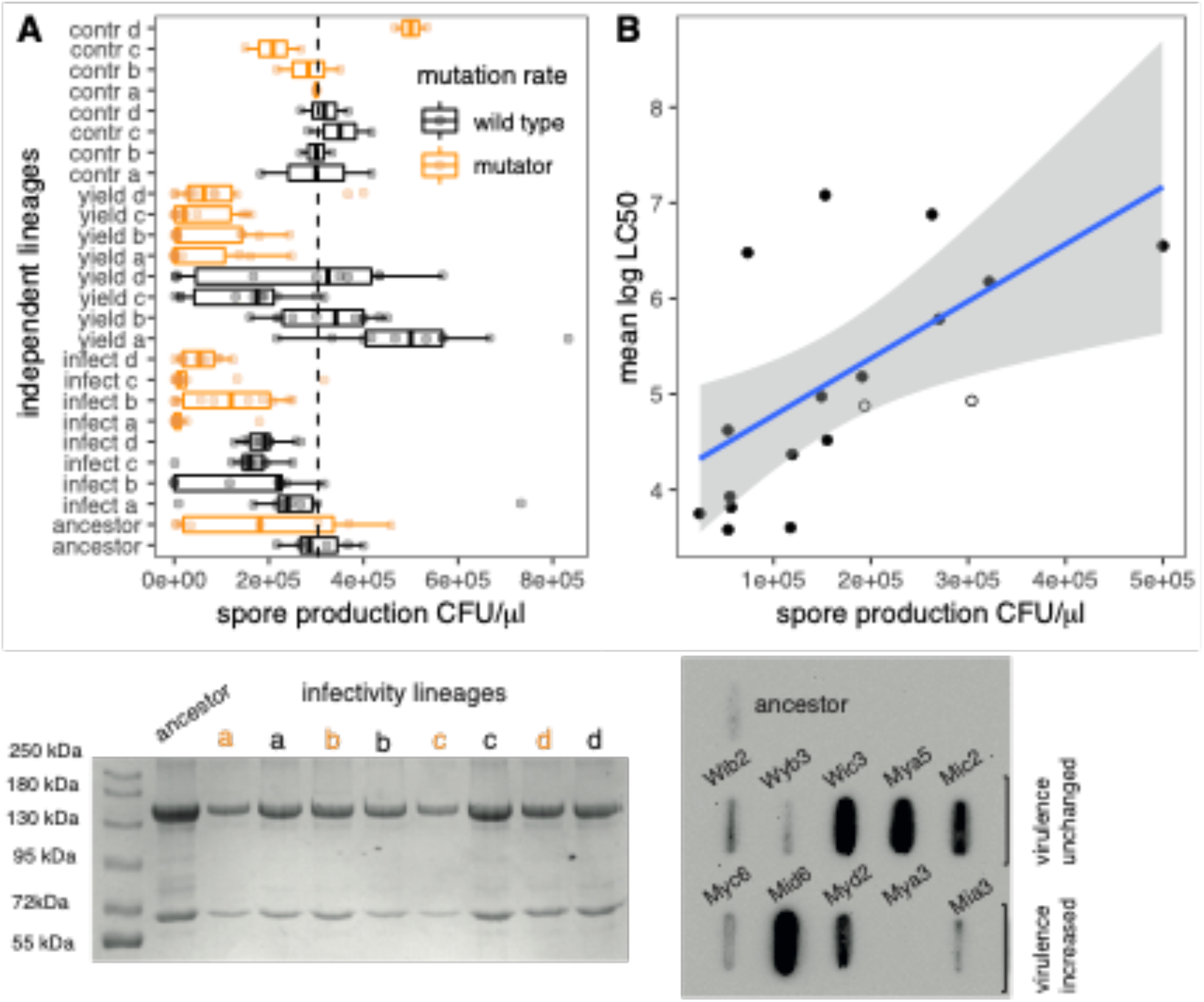
Increased virulence trades-off against reduced spore production in evolved Bt lines but is not associated with increased Vip3 production. (A) Variation in spore production between evolved lineages, asterisks indicate significance of contrasts between all evolved lineages and the ancestor in a *glm*, boxplots summarize the results from two assays of each clone. (B) Mean LC50 of each *in vivo* passaged lineage (averaged across clones) plotted against spore production, ancestors plotted as hollow circles. (C) Clones with increased virulence have delayed sporulation, measured as percentage lysis of cells and release of crystal after 72h growth in sporulation medium; data for *n* = 3 clones and the ancestor. (D) SDS PAGE showing Cry toxin production for all group selected lineages see Fig S3 for examples of individual clones. (E) Slotblot characterization of Vip3 production in the ancestral Bt clone and clones with increased virulence or virulence unchanged relative to the ancestor (see Table S1). See also figures S3, S4, S5 and Tables S2 and S3 for additional details on genomic and proteomic characterization of evolved clones.

Reduced spore production proved to be due to slowed sporulation (Fig 4B), as there were no significant differences in total cell production when cells were grown in sporulation medium. Two classes of toxin are produced by Bt *kurstaki* earlier in the growth cycle than the Cry1A toxins and are expected to have activity against Cry1A resistant insects: these are the Cry2 toxins and the vegetative virulence factor Vip3Aa (Deng et al. 2014; Carrière, Crickmore, & Tabashnik 2015). However, we found no significant differences in the ratio of Cry1A and Cry2 production that correlated with changes in virulence. At the lineage level neither mutators nor infectivity-selected replicates had elevated Cry2 production (Fig 4C), nor did we see differences in Cry2 production between the ancestor and evolved clones with high virulence (Figure S3). We did find substantial variation in Vip3Aa production between evolved clones but no association with increased virulence was seen (Fig 4D).

### Genomic analysis

To seek mechanistic explanations for the reduced sporulation and increased virulence, the genomes of two clones from each lineage were sequenced and compared to a long-read assembly of our ancestor. The ancestral genome completed a 5.7Mb chromosome and 14 plasmids that resolved as single contigs with high similarity to other Bt *kurstaki* genomes (Tables S2-3, Figure S4). Re-sequencing of evolved mutants confirmed that none carried mutations in their Cry genes or in regions immediately upstream. Comparison to the ancestral genome showed both small scale mutations and large-scale deletions associated with plasmid loss. Particularly, toxin gene loss patterns detected previously by PCR can be explained by the loss of one or both Cry plasmids.

Phylogenetic analysis of our evolved clones shows that there was considerably more genome evolution in the mutators (Figure 5A). However, evolved clones with similar virulence phenotypes did not group together, nor did clones from the same treatments (Figure 5A). Mapping of mutations to the ancestral genome also suggests that minimal convergent evolution in terms of shared changes in DNA (although there is phenotypic convergent evolution in terms of sporulation). We did identify a small list of genes that acquired mutations in more than one more than one clone (Figure 5B) this list includes potentially three transcriptional regulators *nprA* (in clones mia3 and mia4) *dagR, sgrR* with non-synonymous mutations in insect passaged lineage, but not in the *in vitro* controls. In general, missense or frameshift mutations were quite common in genes encoding putative transcriptional regulators (supplementary table S4). These included well described regulators such as NprA but also proteins with helix-turn-helix domains. Mutator clones from the infectivity selection treatment accounted for 10 out of the 16 loss of function mutations in putative transcriptional regulators. If we compared the virulence (LC50) of all the sequenced mutator clones there was a significant increase in virulence in the mutator clones with these loss of function mutations compared to those with no mutations in regulatory genes (*F*_1,16_ = 5.91, *P* = 0.027), assuming that the data from these clones are independent observations.

**Figure 5.**
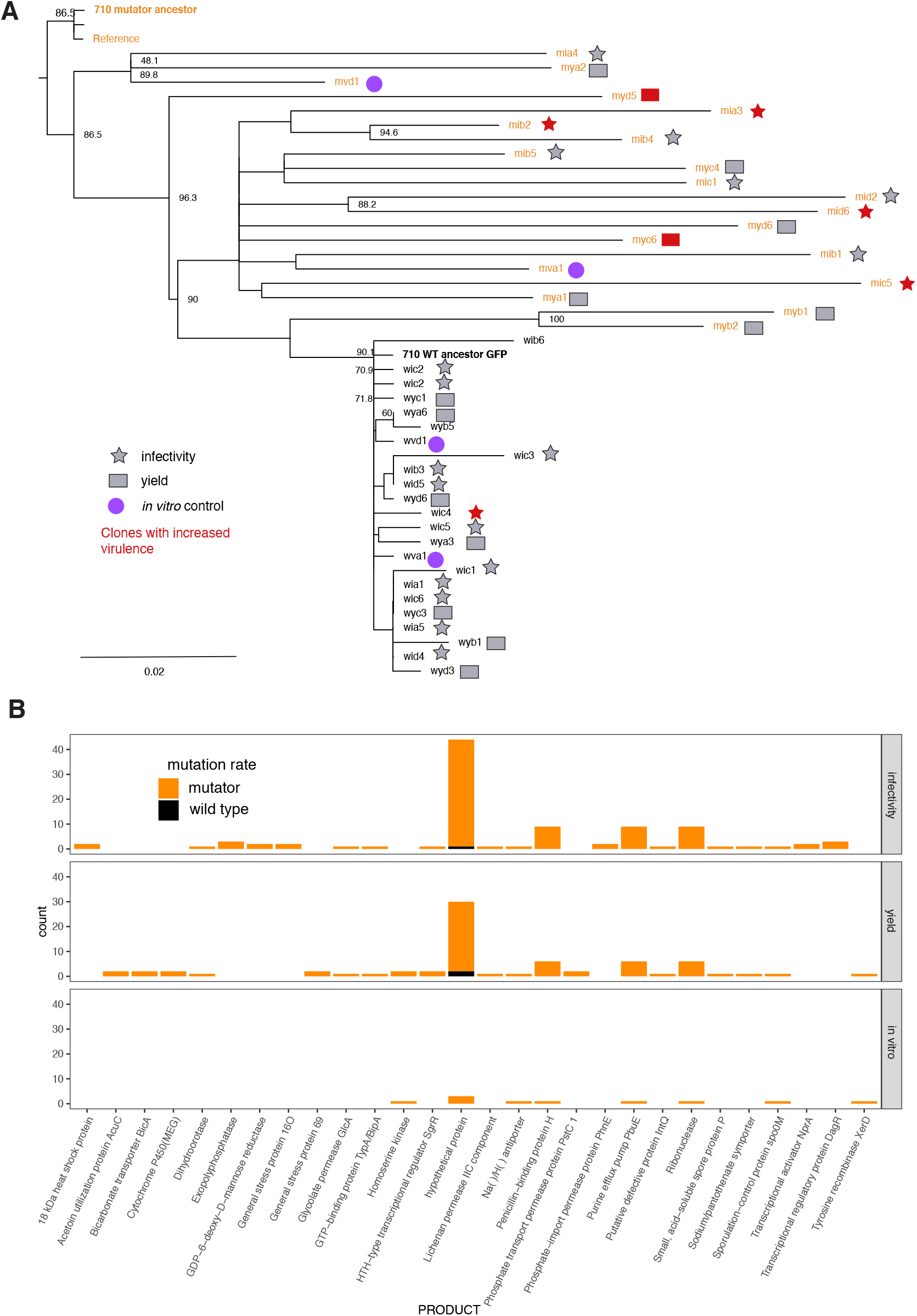
Genomic analyses of evolved Bt clones. (A) consensus Neighbour-joining tree of sequenced evolved clones. For ease of interpretation bootstrap of values > 50 only have been displayed for the nodes. The scale bar indicates number of substitutions per nucleotide. (B). Histogram of non-synonymous mutations identified in at least two evolved clones in our three selection regimes, infectivity selection, yield selection and the *in vitro* passage controls.

**Figure 6.**
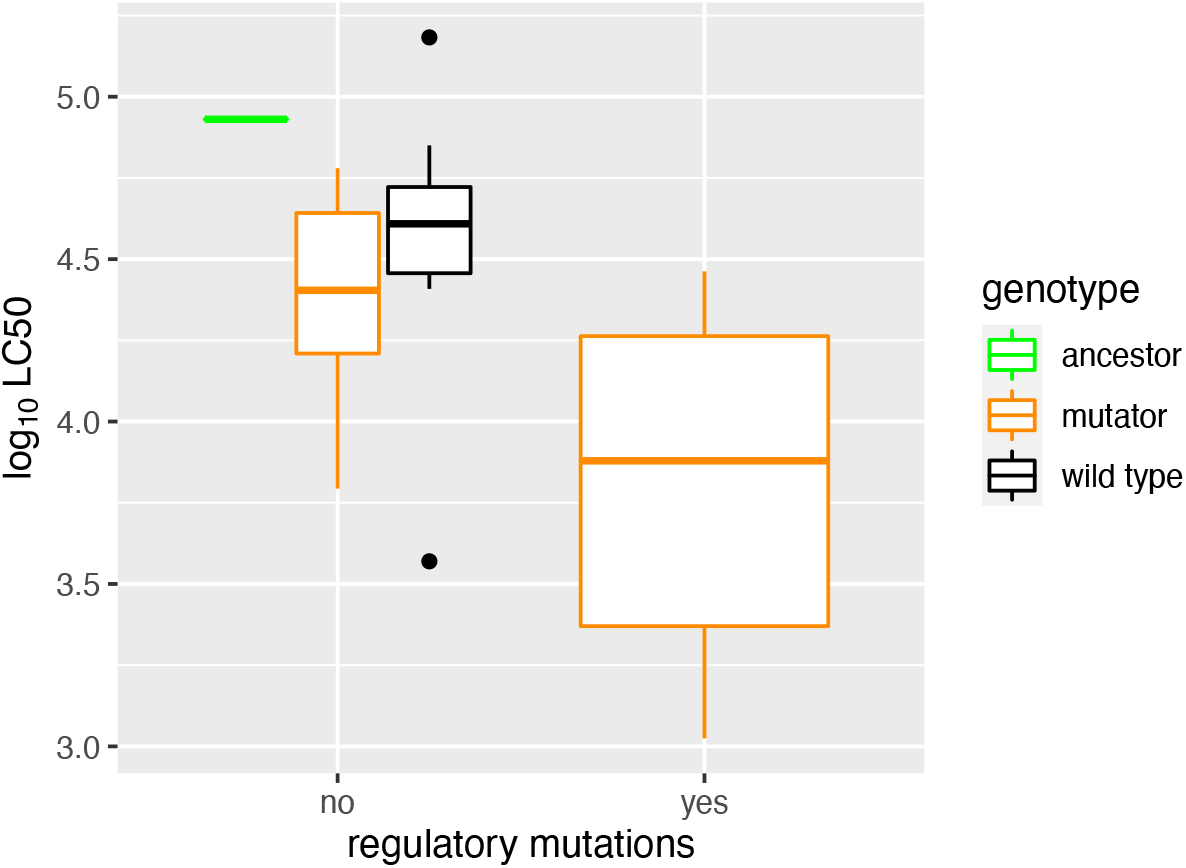
Virulence of evolved mutators clones with and without loss of function mutations in putative transcriptional regulators. Virulence is expressed as log LC50 with evolved wild type clones and the 7.1.o ancestor plotted as comparisons, after excluding data from sequenced clone that are Cry toxin cheaters (*i*.*e*. which carry 0 or 1 Cry toxin genes). Supplementary table S4 contains a list of mutations.

## Discussion

Kin selection theory emphasizes that altered virulence can have different impacts on individual and group level fitness components (Buckling & Brockhurst 2008). Notably investment in bacterial virulence factors requires resources to be diverted away from growth of individual cells. If virulence factors are diffusible, they may behave as ‘public goods’ by conferring fitness benefits that are shared among members of an infecting group (Harrison, Browning, Vos, & Buckling 2006; Raymond et al. 2012; Diard, Sellin, et al. 2014). If cells reduce investment in public goods virulence, these ‘cheaters’ can gain a reproductive advantage during growth in the host by free-loading on the products of others (West & Buckling 2003; Diggle, Griffin, Campbell, & West 2007; Buckling & Brockhurst 2008), although this may be accompanied by a group-level cost such as reduced infectivity. Virulence factors can behave as public goods both in laboratory models and in infections of insects with biocontrol agents such as entomopathogenic nematodes and Bt (Raymond et al. 2012; Zhou et al. 2014; Deng et al. 2015; Shapiro-Ilan & Raymond 2016). In this experiment it was clear that some clones within lineages had reduced virulence relative to the ancestor and so were putative cheaters and we identified a link between the absence of Cry toxins and increases in competitive fitness.

This result mirrors previous experiments with Bt strains that have been cured of Cry toxin encoding plasmids as well as competition between Bt and naturally Cry null *B. cereus* (Raymond, Davis, & Bonsall 2007; Raymond et al. 2012). The key difference in this study is that loss of Cry toxin plasmids occurred spontaneously during experimental evolution in multiple independent lineages and in different ways, *i*.*e*. by the loss of either or both of the large plasmids that encode these toxins. In additional the selection pressure we imposed during experimental evolution affected how readily Cry toxins were lost: selection pressure to maximize yield (final population size in cadavers) resulted in higher rates of plasmid loss than in the infectivity selection treatment. The loss of Cry plasmids in response to selection on yield occurred because Cry toxins impose costs on growth rate *and* total production of spores within hosts (Raymond et al. 2012). Both social evolution and evolution of virulence theory commonly divides the selective forces into individual (within host) and group level (between hosts) components; these are often in conflict in microbes (Frank 1997; Cressler et al. 2016). Other bacterial virulence factors which are public goods, such as siderophores, have a group level benefit in terms increasing yield within the host (West & Buckling 2003; Harrison et al. 2006). However, Cry toxins are produced at sporulation in the cadaver, rather than in the early stages of infection, and these Cry toxins require so much protein that Bt strains which invest in these toxins produce fewer substantially spores per unit resource than their Cry null counterparts (Raymond et al. 2012; Deng et al. 2015). In essence Cry toxins are a different type of public good than siderophores and act to increase infectivity rather than resources available for growth within hosts. It remains to be seen whether other highly bacterial public goods can also reduce the population size of pathogens in hosts but increased infectivity or transmission.

The infectivity selection was not only more effective at retaining Cry plasmids but also resulted in greatest gains in virulence relative to the ancestor. There are several factors that could have contributed to these results. First, budding dispersal in a metapopulation involved less pooling of cadavers than the yield selection treatment. Since individual cadavers are likely to be dominated by different genotypes (van Leeuwen et al. 2015) pooling cadavers can reduce relatedness relativity to infectivity selection treatment. In other words, there is more scope for cheating and greater selection to increase competitive fitness in the yield treatment. An additional consideration is that the infectivity selection also imposed selection for gains in virulence at an additional spatial scale. In all treatments, there was selection for infectivity at a between-host level (successful infections had to occur in order for passage to proceed), but by using selection in a metapopulation we imposed an additional level of competition between subpopulations within a metapopulation. The effects of this level of selection for infectivity in pathogens requires further investigation, but we would hypothesize that it is important when investment in virulence has costs in within host fitness and between host fitness components in terms growth rate and yield of infectious propagules (Raymond & Erdos 2022). It is important to bear in mind that the aim of this study was to devise and test different methods of selection for maintaining and improving virulence and so we will require further work to tease out the precise evolutionary mechanisms behind the success of this particular treatment.

The mutator lines showed clear differences in the rate of molecular evolution as well as greater increases in virulence relative to the wild-type background. In contrast to previous work, we did not see an increase in levels of cheating under high mutation rate (Harrison & Buckling 2005; Racey et al. 2009). Mutagenesis is very commonly employed in strain improvement and directed evolution with *Bacillus* sp and other microbes (Lai et al. 2004; Raju & Divakar 2013). The evidence for the ability of mutators to accelerate increases in fitness is complex and has several unresolved questions. Mutators can increase the supply of both beneficial and deleterious alleles and it is clear there is no known direct benefit to having impaired proof-reading: mutator alleles rise in frequency by hitch-hiking on the fitness benefits of linked alleles (Chao & Cox 1983; de Visser 2002; Gentile et al. 2011; Raynes, Halstead, & Sniegowski 2013). Mutators are often seen at quite high proportions in pathogenic bacteria (LeClerc, Li, Payne, & Cebula 1996; Matic et al. 1997; Oliver et al. 2000) and mutators can appear spontaneously in experimental evolution although evolving lineages often moderate mutation rates during long-term transfer experiments (Good et al. 2017; Ho et al. 2021).

One reason for the prevalence of mutators among pathogens, and for their success in this study is that small populations and / or infection bottlenecks may limit the supply of beneficial mutations (de Visser 2002). While demographics and mutation supply are known to affect the long-term success of mutators, several simulation/experimental studies suggest that bottlenecks or small populations may not favour high mutation rates (Raynes, Wylie, Sniegowski, & Weinreich 2018; Ho et al. 2021). Nevertheless, mutators are more successful when large effect beneficial mutations are available for hitch-hiking (Mao, Lane, Lee, & Miller 1997; Thompson, Desai, & Murray 2006; Gentile et al. 2011) and the strong selection imposed by new sporulation conditions and a novel host genotype in this study may account for the success of mutators in our experiments. Overall, the use of mutators to overcome host resistance has a sensible biological basis. However, this may have had implications for the stability of virulence (Fig S2). Potentially more stable phenotypes can be produced by periodic rounds of chemically induced mutagenesis or by using initial pathogen populations with standing genetic variation.

It did not prove possible to identify a single simple cause for any gains in virulence in this study, although this was consistently related to decreased sporulation efficiency. Reduced sporulation means that vegetative cells and their secreted proteins will be more prevalent in final inocula. We observed that proportion of mutants with reduced sporulation gave increased virulence per spore but not in terms of dilution of the broth in which they were cultured. This is presumably because standardizing doses by spore means that a higher concentration of broth associated virulence factors are included in inocula. As yet, none of the classic virulence factors associated with broth such as Vip3A appear to be responsible for this increase in virulence. There are many virulence factors secreted into broth by Bt and its relatives (Guinebretière, Broussolle, & Nguyen-The 2002; Chitlaru et al. 2006; Stenfors Arnesen, Fagerlund, & Granum 2008). While spores and crystals are by far and away the most significant contributors to virulence in Bt (Raymond et al. 2010), broth associated virulence may be more significant for Cry1A resistant hosts and are known to be significant for some hosts such as *Galleria melonella* (Salamitou et al. 2000).

For example, secreted virulence factors are regulated by a quorum sensing system (PlcR/PapR) that activates a suite of virulence factors involved in host invasion (Salamitou et al. 2000; Zhou et al. 2014). Although these are typically produced at stationary phase, in low nutrient sporulation medium such as HCO expression of the PlcR-regulated genes is repressed (Lereclus et al. 2000). The stationary phase secretome of Bt and *B. anthracis* in sporulation medium is mainly composed of metalloproteases such as Inh1A and NprA and is much reduced in the diversity of proteins compared to nutrient rich media (Chitlaru et al.m 2006; Perchat et al. 2011). These metalloproteases are hypothesized to have a role in the digestion of cadavers in late stage infections (Perchat et al. 2016). Nevertheless, they are essentially lytic enzymes and could improve the ability of Bt to invade the host at high concentration. Other possible virulence factors secreted into sporulating media include chitinase and chitin binding proteins (Chitlaru et al. 2006). Notably we did identify mutations in *NprA* and other transcriptional regulators in several insect-passaged mutators.

Bt is an important pathogen for both organic horticulture and modern ‘biotech’ crops so the question of how to improve strains or discover new toxins is significant, especially given the need to respond to the evolution of resistance (Adang et al. 2014; Badran et al. 2016). However, these methods may be applicable to other parasites, such as nematodes and fungi, which also produce costly excreted virulence factors (Shapiro-Ilan & Raymond 2016). Moreover, in addition to providing a framework for increasing virulence, the combination of *in vivo* and *in vitro* selection used here could be applied when the aim is adaptation to an alternative growth medium or the improvement of other phenotypes, while retaining insecticidal efficacy. For some pathogens, eg fungi, it is also worth bearing in mind that while virulence traits may not be social (there are no data yet), other important biocontrol traits such as sporulation are likely to be subject to social conflicts. The timing of a switch to any resting state (spore, persister etc) can be subject to cheating – *i*.*e*. cessation of growth can be cooperative and late switching to sporulation can provide growth advantages in competition (Gardner, West, & Griffin 2007; Ratcliff, Hoverman, Travisano, & Denison 2013). Efficient sporulation is essential for production of both fungal and bacterial biocontrol agents and can be readily lost during laboratory selection. In general, then passage regimes that minimize social conflict and which can maintain valuable cooperative traits could have broad importance across diverse groups of invertebrate pathogens, and here we have shown the value of a metapopulation regime that can reduce social conflicts. Moreover, experimental evolution offers a mechanism-free solution to overcoming pest resistance is potentially applicable to many pathogens and pests, and may help us discover entirely novel virulence factors or virulence modification responses in contrast to directed evolution methods that require well-understood protein-protein interactions (Badran et al. 2016).

## Data Archiving statement

Sequence data are hosted at the SRA under BIOPROJECT SUB10359598. Plasmids, ancestor and mutator clones are available on request. Evolved mutants can be shared subject to Material Transfer Agreements. Experimental data and statistical code will be available via a permanent doi at our institutional repository (Open Research Exeter) after acceptance in a peer reviewed journal.

## Acknowledgments

thanks to Andy Matthews for support and technical advice. This study was supported by the Leverhulme Trust (RPG-2014-252) and BBSRC (BB/S002928/1)

## Supplementary Materials

Figure S1. Variation in spore associated virulence across evolved *in vivo* clones

Figure S2. Comparison of the virulence of 5^th^ round evolved whole populations and clonal isolates

Figure S3. SDS PAGE gels showing the relative production of Cry1 and Cry2 toxins

Figure S4. Comparison of plasmid genomes between the reference strain HD1 and the 71o strain (ancestor for this study).

Table S1. Virulence (LC50) of selected clones from the passage experiment.

Table S2. Summary of plasmid comparison between reference strain HD1 and the 71o strain.

Table S3. Coverage of 71o strain short read sequence data mapping to the HD1 reference strain.

Table S4. Passaged clones with mutations in putative transcriptional regulators.

