## Supplemental figures and tables for "Re-inventing pathogen passage for social microbes"

**Figure S1. Variation in spore associated virulence across evolved *in vivo* clones.** Logit-transformed mortality 5 days after inoculation is shown as a function of Bt dose as $\log_{10}$ heat tolerant CFU per μl (**A**) and as a volume of standard diluted inoculant instead of dose (**B**), to avoid false positives arising from low plating efficiency/very low sporulation for some clones The black line in each panel describes the dose response for the ancestor (from a glm with quasibinomial errors).

**A**

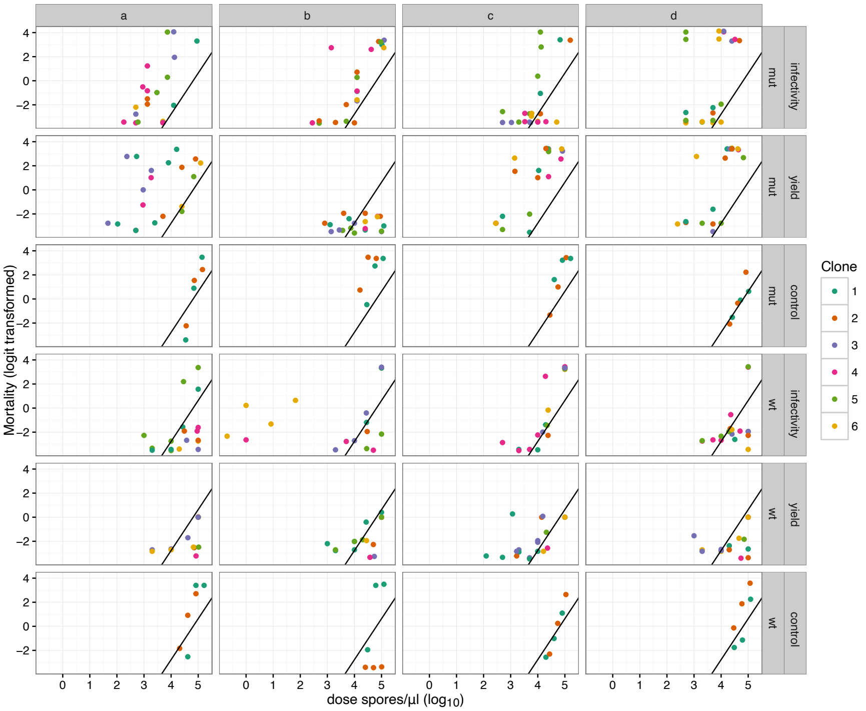


**B**

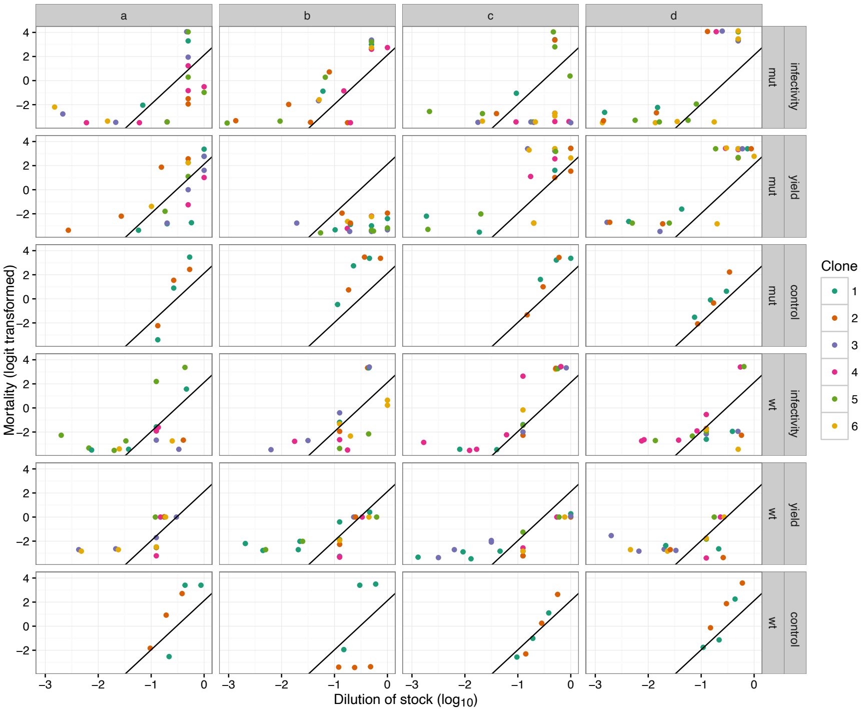

**Figure S2. Comparison of the virulence of 5th round evolved whole populations and clonal isolates.** Logit-transformed mortality 5 days after inoculation is shown as a function of Bt dose. Bt dose is shown as volume of standard diluted inoculant instead of dose (CFUs), to avoid false positives arising from low plating efficiency for some clones. Clone data outside of the range of doses tested for population assays were excluded. Virulence of mutator (mut) populations was significantly higher than that of mut clones, whereas virulence of wild type (wt) populations was not different from that of wild type clones (post-hoc Tukey tests on logit-transformed mortality ~ dose * treatment * clone:strain, mut mix > mut clones, z=10.9, p<0.001; wt mix < wt clones, z=-0.93, p=0.78) suggesting that mutators had reduced phenotypic stability relative to wt clones.

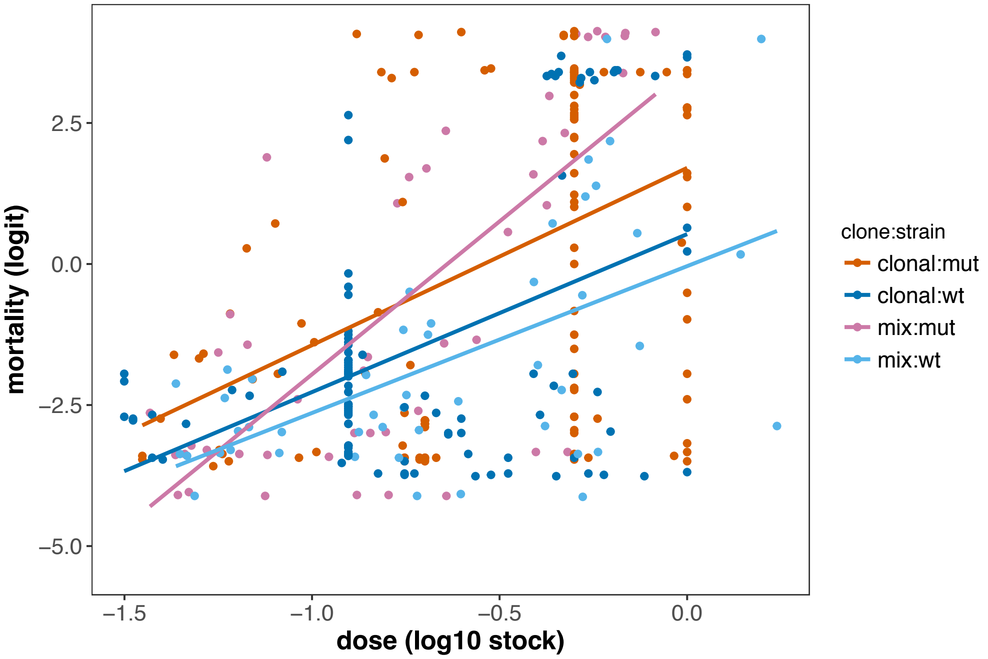


**Figure S3. SDS PAGE gels showing the relative production of Cry1 and Cry2 toxins (at approximately 140 and 70 kDa respectively) by the Bt *kurstaki* ancestor and two evolved clones**. Mia3 and Myc6 both showed increases in virulence relative to the ancestor (see Table S1). Culture methods replicated those in the selection experiment (three days in HCO); equal culture volumes were loaded in this gel.

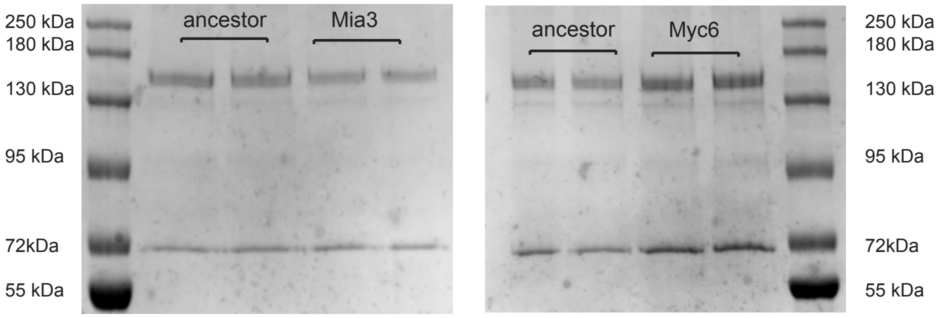

**Figure S4. Comparison of plasmid genomes between the reference strain HD1 and the 71o strain (ancestor for this study).** The graphic is to scale and based on a Mauve (https://doi.org/10.1371/journal.pone.0011147) output annotated to better show inverted, unique regions and missing plasmids between the two strains. The blocks of colour and the grey joining ribbons correspond to homologous regions between the strain plasmids. Plasmid numbers are matched to reference accession in Table S2**.**

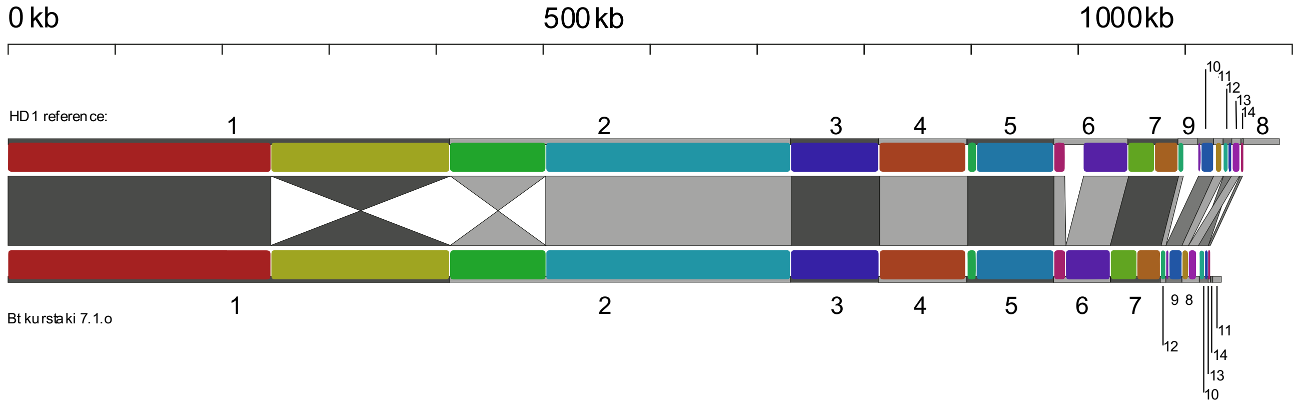

**Table S1 Virulence (LC50) of selected clones from the passage experiment.** LC50 concentrations are in $\log_{10}$ CFU $\mu L^{-1}$. Changes in virulence with respect to ancestor were first tested in clone level analyses of all bioassay data using glms and quasibinomial doses with clone and $\log_{10}$ CFU $\mu l^{-1}$ fitted as main effects. A subset of clones with putative increased virulence were re-assayed; differences in virulence with respect to the ancestor in post-hoc treatment contrasts are identified in this table (Significance of tests: * $p < 0.05$, ** $p < 0.01$; *** $p < 0.001$). For a subset of clones with increased virulence we present LC50s and their standard errors calculated using the *dose.p* function using the *MASS* package after fitting simple logit models using $\log_{10}$ dose and quasibinomial errors. We also list additional clones used in the proteomic analysis in Figure 4 C, D & E which did not increase virulence during passage: NS indicates a LC50 that is not significantly different from the ancestor. †These clones have also been assayed in VLSS insects (Bt susceptible) and show no loss of virulence in that host.

| *Clone* | *Genetic Background* | *$Log_{10}$ LC50* (SE). NA indicates indistinguishable from ancestor | *Increase in virulence relative to ancestor* |
|---|---|---|---|
| ancestor | wild-type | 5.00 (0.1) | - |
| Mia3 | mutator | 3.82 (0.13) | 15.1 *** |
| Mib2 | mutator | 4.21 (0.08) | 6.2 *** |
| Mic5 | mutator | 3.99 (0.4) | 10.2 *** |
| Mid6† | mutator | 3.69 (0.3) | 20.4 *** |
| Mya3 | mutator | 4.4 (0.2) | 3.1 * |
| Myc6 | mutator | 3.9 (13.2) | 11.2 ** |
| Myd5† | mutator | 4.24 (0.36) | 5.8 ** |
| Wic4† | wild type | 4.44 (0.09) | 3.6 *** |
| Mic2 | mutator | NA | NS $p = 0.57$ |
| Mya5 | mutator | NA | NS $p = 0.44$ |
| Wib2 | wild-type | NA | NS $p = 0.76$ |
| Wic3 | wild-type | NA | NS $p = 0.47$ |
| Wyb3 | wild-type | NA | NS $p = 0.33$ |

**Table S2. Summary of plasmid comparison between reference strain HD1 and the 71o strain.**

| Current study: strain 71o | | | | HD1 reference | | | |
|---|---|---|---|---|---|---|---|
| **Contig** | **Plasmid no.** | **Length (bp)** | **Depth relative to chromosome** | **Plasmid no.** | **HD1 accession** | **Length (bp)** | **Size difference (bp)** |
| Chromosome 1 | NA | 5,655,593 | 1.00x | na | CP010005 | 5,675,189 | (19,596) |
| 2 | 1 | 412,028 | 2.08x | 1 | CP009998 | 412,022 | 6 |
| 3 | 2 | 317,339 | 4.43x | 2 | CP009999 | 317,336 | 3 |
| 4 | 3 | 82,377 | 2.21x | 3 | CP010000 | 82,528 | (151) |
| 5 | 4 | 82,304 | 2.88x | 4 | CP010001 | 82,303 | 1 |
| 6 | 5 | 80,699 | 1.15x | 5 | CP010002 | 80,285 | 414 |
| 7 | 6 | 52,548 | 4.20x | 6 | CP010003 | 69,317 | (16,769) |
| 8 | 7 | 46,634 | 10.19x | 7 | CP010006 | 46,634 | - |
| 9 | 8 | 16,148 | 26.38x | 12 | CP010007 | 8,513 | 7,635 |
| 10 | 9 | 14,889 | 4.65x | 10 | CP010010 | 14,889 | - |
| 11 | 10 | 8,279 | 10.69x | 11, 13 | CP010009, CP010011 | 15,914 | (7,635) |
| 12 | 11 | 7,705 | 17.77x | no match | no match | | 7,705 |
| 13 | 12 | 5,113 | 14.80x | 9 | CP010012 | 18,332 | (13,219) |
| 14 | 13 | 2,062 | 23.33x | 14 | CP010008 | 2,062 | - |
| 15 | 14 | 1,944 | 11.34x | no match | no match | | 1,944 |
| Total (bp) | | 6,785,662 | | | | 6,825,324 | 73812 |

**Table S3. Coverage of 71o strain short read sequence data mapping to the HD1 reference strain**. Confirming the absence of HD1 plasmid CP010004 in the 71o genome.

| HD1 accession | Plasmid number | HD1 reference size | number of short reads mapping | coverage | copy number vs chromosome |
|---|---|---|---|---|---|
| CP010005 | Chromosome | 5,675,189 | 3,054,392 | 48 | 1.0 |
| CP009998 | 1 | 412,022 | 442,079 | 97 | 2.0 |
| CP009999 | 2 | 317,336 | 22,100 | 205 | 4.2 |
| CP010000 | 3 | 82,528 | 111,588 | 122 | 2.5 |
| CP010001 | 4 | 82,303 | 132,927 | 145 | 3.0 |
| CP010002 | 5 | 80,285 | 118,666 | 133 | 2.7 |
| CP010003 | 6 | 69,317 | 166,436 | 216 | 4.5 |
| CP010006 | 7 | 46,634 | 270,602 | 522 | 10.8 |
| CP010004 | 8 | 34,150 | | | |
| CP010012 | 9 | 18,331 | 208,629 | 1,024 | 21.1 |
| CP010010 | 10 | 14,889 | 39,498 | 239 | 4.9 |
| CP010007 | 11 | 8,513 | 152,832 | 1,616 | 33.4 |
| CP010009 | 12 | 8,279 | 62,718 | 682 | 14.1 |
| CP010011 | 13 | 7,635 | 78,095 | 921 | 19.0 |
| CP010008 | 14 | 2,062 | 27,347 | 1,194 | 24.6 |

**Table S4. Passaged clones with mutations in putative transcriptional regulators.** HTH denotes Helix turn Helix proteins.

| Background | Selection treatment | Lineage | Clone | Protein | Type of mutation |
|---|---|---|---|---|---|
| Mutator | Infectivity | mga | 3 | NprA | stop gained |
| Mutator | Infectivity | mga | 4 | NprA | frameshift |
| Mutator | Infectivity | mgb | 2 | DagR | missense |
| Mutator | Infectivity | mgb | 4 | DagR | missense |
| Mutator | Infectivity | mgb | 4 | repressor PagR | missense |
| Mutator | Infectivity | mgc | 1 | DagR | missense |
| Mutator | Infectivity | mgc | 1 | LiaR | missense |
| Mutator | Infectivity | mgc | 5 | HTH-type regulator SgrR | missense |
| Mutator | Infectivity | mgc | 5 | HTH-type repressor KstR2 | missense |
| Mutator | Infectivity | mgd | 2 | HTH-type regulator | frameshift |
| Mutator | Infectivity | mgd | 2 | activator HcaR | frameshift |
| Mutator | Yield | mpa | 1 | HTH regulator ImmR | missense |
| Mutator | Yield | mpa | 2 | LiaR | missense |
| Mutator | Yield | mpa | 2 | HTH-type regulator SgrR | frameshift |
| Mutator | Yield | mpc | 6 | HTH-regulator LutR | frameshift |
| Mutator | Yield | mpd | 6 | HTH-regulator NorG | missense |
| Mutator | *in vitro* | mva | 1 | HTH-regulator CysL | missense |
| | | Synonymous mutations | | | |
| Mutator | *in vitro* | mvd | 1 | HTH-regulator CysL | synonymous |
| Mutator | *in vitro* | mvd | 1 | HTH-regulator CatM | synonymous |
| Mutator | Infectivity | mga | 4 | HTH-repressor ComR | synonymous |
